# A co-opted endogenous retroviral envelope promotes cell survival by controlling SLC31A1/CTR1-mediated copper transport and homeostasis

**DOI:** 10.1101/2021.12.09.471612

**Authors:** Sandrine Tury, Lise Chauveau, Valérie Courgnaud, Jean-Luc Battini

## Abstract

Copper is a critical element for eukaryotic life, involved in numerous cellular functions and in redox balance but it can be toxic in excess. Therefore, tight regulation of copper acquisition and homeostasis is essential for cell physiology and survival. Here, we identified a unique mechanism for cell survival involving the regulation of copper homeostasis by an endogenous retroviral (ERV) envelope glycoprotein called Refrex1. We show that extracellular copper sensing by cells increases Refrex1 expression, which in turn regulates copper acquisition through interaction with the main copper transporter SLC31A1/CTR1. Downmodulation of Refrex1 resulted in intracellular copper accumulation leading to ROS production and subsequent apoptosis, which could be reverted by copper chelator treatment. Our results demonstrate that Refrex1 has been co-opted for its ability to regulate copper entry through CTR1 interaction in order to limit copper excess for a proper redox balance, and suggests that other ERV may have similar metabolic functions among vertebrates.

## INTRODUCTION

Copper is an unmissable element for the development of all forms of eucaryotes. Copper ions exist in two redox states, the cuprous ion Cu^+^ and the cupric ion Cu^2+^. This Cu^+^/Cu^2+^ redox couple is a powerful cargo of electrons and is therefore critical for a multitude of enzymatic reactions that are essential for cellular functions, including cellular respiration and free radical detoxification^1,2^. However, copper excess is toxic as it affects the cellular redox balance leading to unwanted toxic reactive oxygen species (ROS). Therefore, copper homeostasis is tightly controlled to meet both copper sufficiency and toxic limitation. It relies on regulated mechanisms involving multiple cellular factors with key roles in copper acquisition, in sequestering and chaperoning copper for proper distribution, and in excretion. Copper transporters are particularly important in this process, with the ATPase pumps ATP7A and ATP7B concentrating copper into the trans-Golgi network and the main plasma membrane-localized copper transporter 1 SLC31A1/CTR1 importing Cu^+^ from the extracellular environment^2^. In situation of copper excess, ATP7A and ATP7B relocate to the plasma membrane of human cells and release copper while CTR1 is endocytosed to prevent excessive copper uptake^3^. Other cellular adaptations include the cleavage of the CTR1 ectodomain by cathepsins in a CTR2-dependent manner leading to reduced copper binding to CTR1 and subsequent uptake^4–6^. Such specialized mechanisms have been adopted by organisms during evolution which suggests that other important regulators of the copper homeostasis machinery may exist in vertebrates.

Retrovirus-vertebrate interaction is a recognized factor of evolution which has shaped genomes with hundreds of inherited retroviral sequences^7^. In humans, 42% of genomic DNA sequences have retroviral origins, of which 8% are endogenous retroviruses (ERV) with a genetic organization similar to modern exogenous retroviruses^8,9^. Although most ERV are inactive due to accumulation of deleterious mutations^9^, rare ERV genes have been co-opted by hosts for cellular functions. To date, co-option of more than 80 retroviral *env* genes have been identified in vertebrates^10^, including 18 in humans^11,12^. These *env* encode envelope glycoproteins (Env) with receptor recognition, membrane fusion and immunosuppressive properties^13^. Co-opted Env retaining receptor-binding properties have often been described as restriction factors in various animals for their capacity to confer resistance to exogenous infections *via* receptor saturation ^14–18^, but several of them have been reassigned by hosts for physiological functions. This includes the human ERV, HERV-W and HERV-FRD Env, renamed syncytins for their role in syncytiotrophoblast formation during placentation^19,20^, as well as non-human syncytins found in numerous mammals^13^. Also, a co-opted Env originating from HERV-K subtype HML2 was shown to be involved in stemness maintenance to avoid neuronal differentiation through the regulation of the mTOR pathway^21^. Most ERV originated from ancient gamma-retroviruses and all their retroviral receptors identified so far are members of a large family of cell surface nutrient transporters involved in cell metabolism^22^, thus their interaction with retroviral Env often affects nutrient transport^23–26^. Therefore, in addition to their role as restriction factors, as regulator of Stemness or as fusogenic molecules for placentation, it is not excluded that endogenous Env could also act as modulators of cellular metabolism.

In domestic cats, two endogenous categories of ERV express truncated receptor-binding domain (RBD) Env, namely FeLIX and Refrex1^16,27^. FeLIX is related to the feline leukemia virus type B (FeLV-B) endogenous Env^28^ and as such is able to bind the SLC20A1/PiT1 phosphate transporter^29^. On the other hand, Refrex1 is related to ERV-DC^30^, is expressed from the ERV-DC7 and ERV-DC16 loci, and has been described as a restriction factor against ERV-DC infection^16,31^. ERV-DC7 and 16 loci are fixed in domestic cats as well as in European wild cats (ERV-DC7 only) and are actively transcribed, suggesting that Refrex1 activity has been evolutionary conserved in cats^31–33^. In this study, we asked whether Refrex1, in addition to its role as a restriction factor, has been co-opted for a more physiological function. We report here that Refrex1 is involved in cell survival by controlling copper homeostasis and redox balance through interaction with the copper transporter CTR1. Thus, co-option of endogenous *env* genes by the host could also be envisioned as a selective advantage for proper cell metabolism and survival.

## RESULTS

### ERV-DC7/16 *env* gene silencing in feline FEA cells

Refrex1 was first described as a soluble antiretroviral factor present in wild and domestic cats and in the culture medium of several cat cell lines^16^, but it is not known whether Refrex1 also plays a physiological role in these cells. We first evaluated by RT-qPCR the expression level of the 3 genotype groups of ERV-DC *env* transcripts in feline FEA cells, namely group I (ERV-DC2, 8, 14, 17, 19), group II (ERV-DC7, 16) encoding Refrex1, and group III (ERV-DC6, 10, 18). We found that group II mRNA were highly expressed in FEA cells in much higher amount than those of groups I and III (Fig 1A, left panel), suggesting a predominance of Refrex1 expression over the other ERV-DC Env. Surprisingly, we only detected the expression of the ERV-DC16 mRNA in FEA cells, and the same observation was made in CRFK cells, suggesting that the ERV-DC7 locus was inactive in these 2 feline cell lines (Supplementary Fig 1). Nevertheless, both ERV-DC7 and 16 mRNA were expressed in cat PBMCs lines (Supplementary Fig 1). We next designed a siRNA (siDC7/16) targeting a common sequence in ERV-DC7 and ERV-DC16 *env* genes in order to silence Refrex1 expression (Supplementary Fig 2). Transfection of FEA cells by siDC7/16 but not by a siRNA control targeting the *firefly luciferase* gene (siLuc) decreased the level of group II mRNAs significantly and to a lesser extent those of group I despite 2 mismatches within the siDC7/16 sequence (Fig 1A, right panels). All feline cell lines are resistant to ERV-DC14 pseudotype infection, including FEA cells (Fig. 1B). We observed that Refrex1 silencing increased by 20-fold the infectivity of lentiviral vector particles carrying the *GFP* gene pseudotyped with the ERV-DC14 Env (Fig. 1B), and increased the binding of an ERV-DC14 immunoadhesin comprising a Env RBD fused to a mouse IgG1 Fc fragment (DC14RBD) (Fig. 1C). In contrast, binding of the FLVC Env RBD (FLVCRBD) which recognizes the heme transporter FLVCR1 was unchanged. Thus, presence of Refrex1 in FEA cells is responsible for the resistance to ERV-DC14 pseudotype infection. We also tested the presence of Refrex1 in FEA-conditioned medium (CM) by measuring its inhibitory activity on ERV-DC14 infectivity and binding. Pretreatment of 293T cells with FEA-CM demonstrated a strong decrease in both ERV-DC14 pseudotype infection as expected^16^ and DC14RBD binding, while infectivity of VSVG-pseudotyped particles was unchanged as well as FLVCRBD binding (Fig 1D-E). In contrast, CM of siDC7/16-transfected FEA cells was unable to inhibit Env binding and infection with ERV-DC14 (Fig 1D-E), suggesting that the release of Refrex1 in FEA CM was abolished. This also demonstrated that Refrex1 was solely expressed from ERV-DC7 and 16. Unexpectedly, we found that Refrex1 silencing was accompanied by cell toxicity as observed by the low percentage of gated cells in the siDC7/16 dotplot of Fig. 1C.

**Figure 1.**
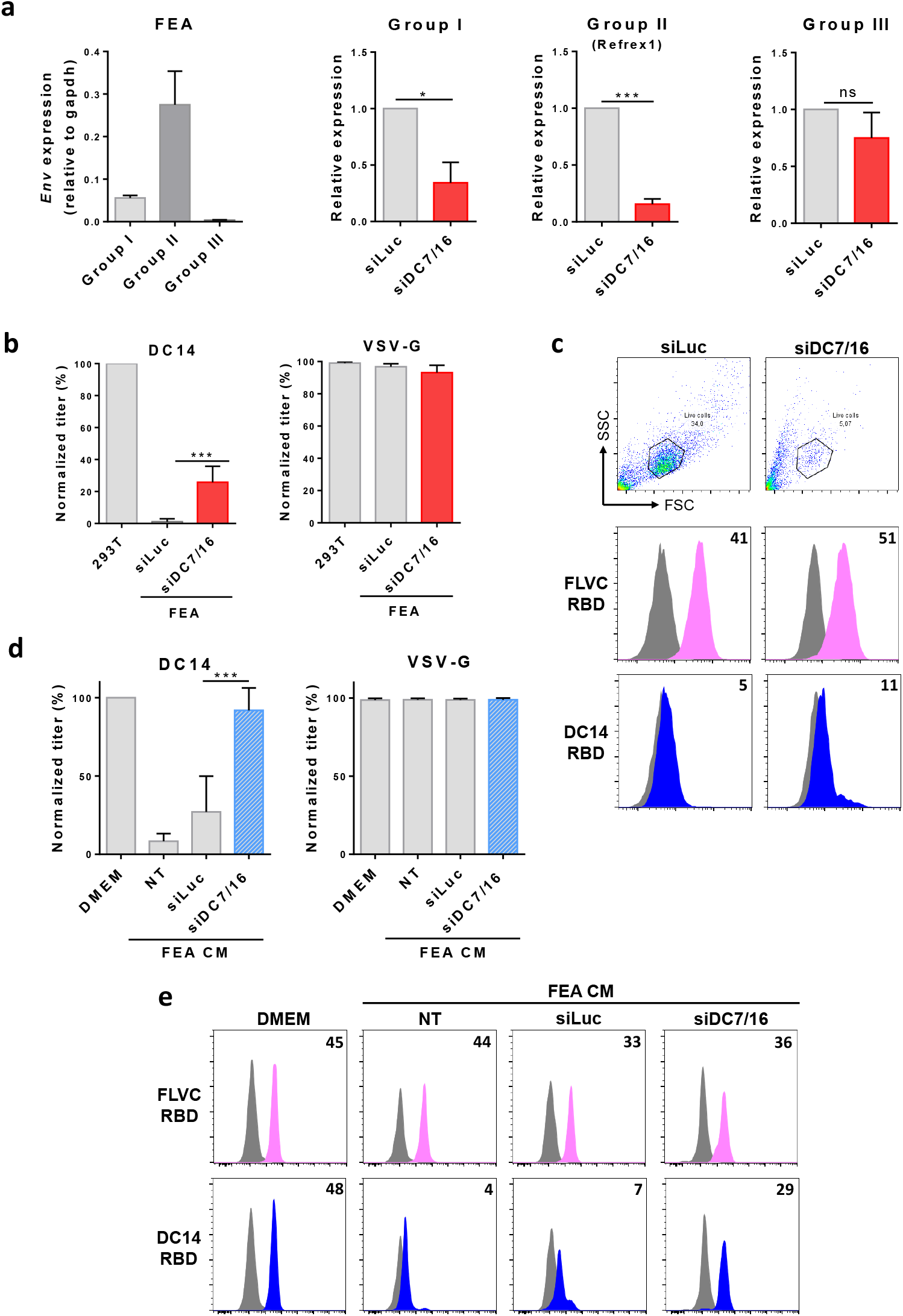
ERV-DC7/16 *env* gene silencing in feline FEA cells. **(a)** ERV-DC *env* gene expression from genotype groups I, II, or III was evaluated by RT-qPCR cells and normalized with cat Gapdh in non-transfected FEA cells (left panel), or after transfection with siDC7/16 targeting both ERV-DC7 and DC16 *env* or with a siRNA control (siLuc) (3 right panels). Fold changes are normalized to siLuc. Data are means ± SEM from n=3 experiments. Student unpaired t-test * = p ≤ 0.05 *** = p ≤ 0.001 **(b)** Sensitivity of siDC7/16 or siLuc-transfected FEA cells to infection by EGFP lentiviral vectors pseudotyped with ERV-DC14 or VSV-G Env. 293T target cells were used as control and for normalization. Data are means ± SEM from n=3 experiments. Student unpaired t-test *** = p ≤ 0.001 **(c)** Cells from (b) were evaluated for FLVC and DC14 Env RBD binding by flow cytometry. Forward scatter (FSC) *versus* Side scatter (SSC) plots show gated living cells. Numbers indicate the specific change in mean fluorescence intensity compared to mock (grey histogram). Representative experiment of n=3. **(d)** 293T incubated in conditioned medium (CM) from feline FEA cells transfected or not with the indicated siRNA or in DMEM for 5h were evaluated for their sensitivity to infection by EGFP lentiviral vectors pseudotyped with VSV-G or ERV-DC14 Env. NT: non transfected. Data are means ± SEM from n=3 experiments. Student unpaired t-test *** = p ≤ 0.001. **(e)** Cells from (d) were evaluated for RBD binding as in (c). Representative experiment from n=3.

### Loss of Refrex1 expression induces cell death by apoptosis

The concomitant loss of Refrex1 and appearance of cell toxicity upon siDC7/16 transfection may suggest a role of Refrex1 in cell survival. Transfection of FEA cells by 2 different siRNA able to silence Refrex1, siDC7/16 and siDC16 targeting only the DC16 *env* gene (Supplementary Fig 2), confirmed the observed toxic effect as seen by phase-contrast microscopy and by enzymatic assay (Fig 2A-B), at levels similar to glutathione peroxidase 4 (GPX4) silencing known to induce ferroptosis^34^. Toxicity was not specific to FEA cells since we observed the same effect in CRFK cells, albeit less pronounced, and this in a dose dependent manner (Supplementary Fig 3). Moreover, toxicity was readily observed 24 hours after transfection in FEA cells, but took longer time in CRFK cells, suggesting that both cell lines have different sensitivity toward Refrex1 depletion (Supplementary Fig 3). In contrast, this defect was observed neither with the siLuc control nor with siDC7/16 and siDC16-transfected human cells that do not express Refrex1 (Supplementary Fig 4). To assess whether this was due to an indirect effect or was attributed specifically to DC16 Env loss, we performed a rescue experiment assay using a DC16 *env* expression vector. As expected, transfection of FEA and CRFK cell lines with siDC16 affected cell survival even in combination with an empty expression vector (Fig 2C). In contrast, cell survival was fully restored after re-expression of DC16 Env (Fig 2C) from an expression vector insensitive to siDC16 (Supplementary Fig 2). We next asked what type of cell death was responsible for the observed cell toxicity. Compared to mock-transfected FEA and CRFK cells, Refrex1 silencing induced a cell death characterized by an increase in the percentage of annexin V-positive cells at 24h post-transfection, a time point before excessive cell death (Fig 2 D). The cell death was delayed in CRFK cells that showed a lower proportion of annexin V-positive cells at 24h (Fig 2 D). Thus, these results strongly point to Refrex1 as an essential factor protecting feline cells from apoptosis.

**Figure 2.**
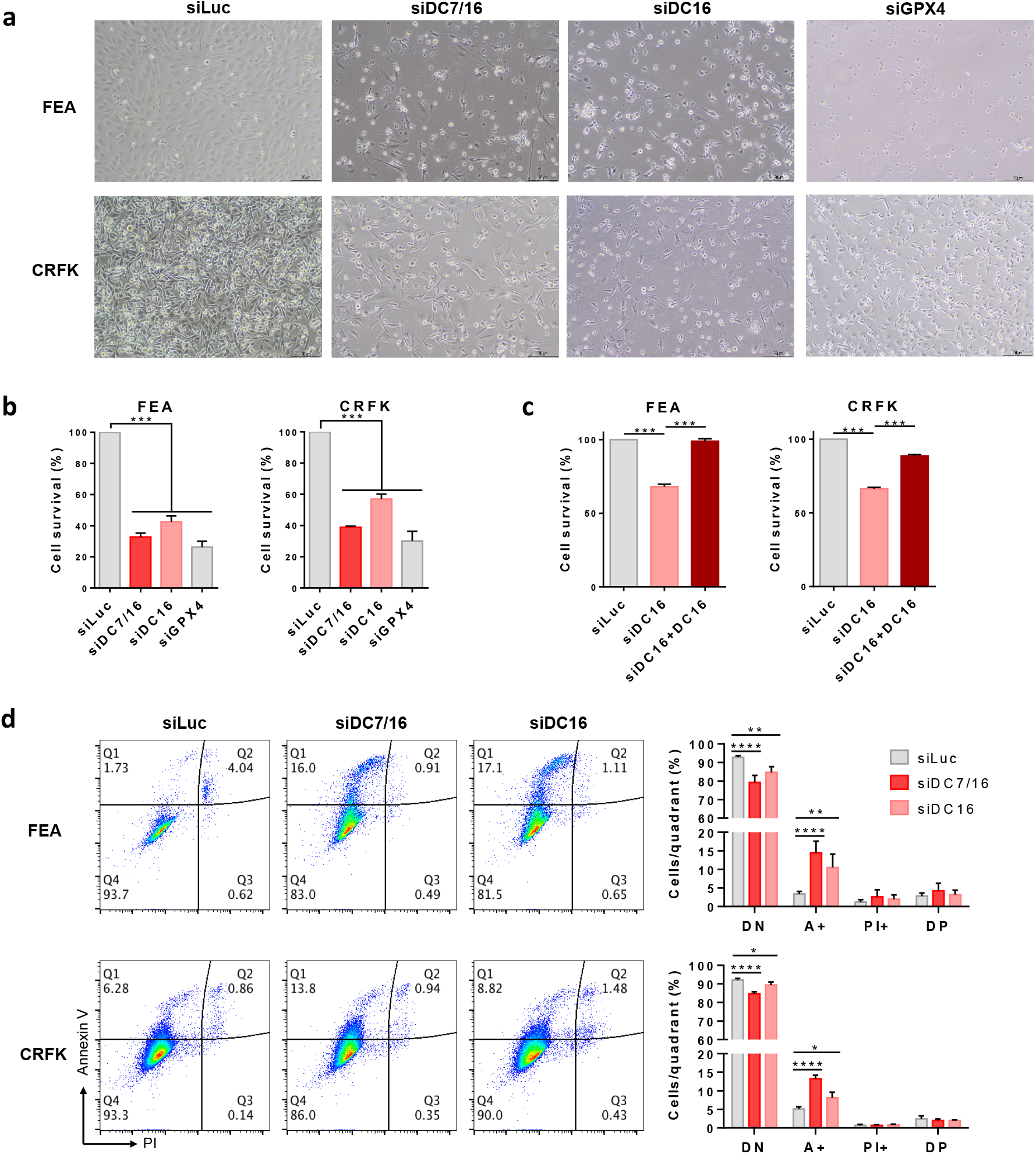
Inactivation of Refrex1 expression induces cell death. **(a)** Representative images of feline FEA and CRFK transfected with either siDC7/16, siDC16, siLuc control or a siRNA targeting a siRNA targeting feline GPX4 mRNA (siGPX4) under bright light using Nikon Eclipse Ts2 microscope. Scale bars = 70 µm **(b)** Cellular viability of cells from (a) was evaluated 48h post-transfection using CellTiter 96® AQueous One Solution Cell Proliferation Assay. Data are means ± SEM from n=3 experiments. One-way ANOVA with Dunnett’s multiple comparisons test *** = p ≤ 0.001. **(c)** FEA and CRFK cells were transfected with siLuc, siDC16 with an empty vector or with DC16 env expression vector. Cell viability was evaluated as in B. Data are means ± SEM from n=3 experiments. One-way ANOVA with Dunnett’s multiple comparisons test *** = p ≤ 0.001. **(d)** FEA and CRFK transfected with siLuc, siDC7/16 or siDC16 were stained with Annexin V/PI at 24h post transfection. Pseudocolour plots from a representative experiment (left panels) and means ± SEM from at least n=3 experiments (right histograms) are shown. DN = double negative cells, A+ = Annexin V positive cells, DP = double positive cells. Two-way ANOVA with Sidak’s multiple comparisons test* = p ≤ 0.05, ** = p ≤ 0.01, **** = p ≤ 0.0001.

### Refrex1 interacts with CTR1

Since Refrex1 is an Env ligand able to interfere with ERV-DC14 infection, we reasoned that its interaction with a specific cell surface receptor would be a critical step to prevent cell death. We previously identified the copper transporter CTR1 as a retroviral receptor for ERV-DC14 infection (Fig 3A)^35^. We therefore tested the possibility that Refrex1 could bind CTR1 at the plasma membrane. We first generated a Refrex1 ligand by introducing 3 copies of the HA tag in place of the stop codon of ERV-DC16 *env* (DC16HA). DC16HA elicited strong binding on human 293T cells but no binding was observed on hamster CHO cells (Supplementary Fig 5). Introduction of HA-tagged human or feline *CTR1* cDNA into hamster CHO cells (Fig. 3B) facilitated the binding of DC16HA but did not alter the binding of the FLVCRBD (Fig. 3C). Therefore, our results suggest that CTR1 is a potent and specific receptor for Refrex1. We next asked whether CTR1 was the only cell surface receptor for Refrex1. Inactivation of the *CTR1* gene in 293T cells by genome editing fully abolished the global expression of CTR1 (Fig. 3D) as well as its specific presence at the plasma membrane (Fig. 3E) using an anti-human CTR1 antibody. Depletion of CTR1 led to a dramatic loss of DC16HA binding while FLVCRBD binding remained unchanged (Fig. 3E). Another mean of downmodulating CTR1 expression from the surface of human cells is exposure to high dose of copper^3,36^. Fig. 3F indicated that overnight treatment of human 293T cells with 100 µM of copper chloride specifically decreased DC16HA binding, while magnesium chloride had no effect. Overall, these data indicated that CTR1 is a specific and the sole receptor for Refrex1.

**Figure 3.**
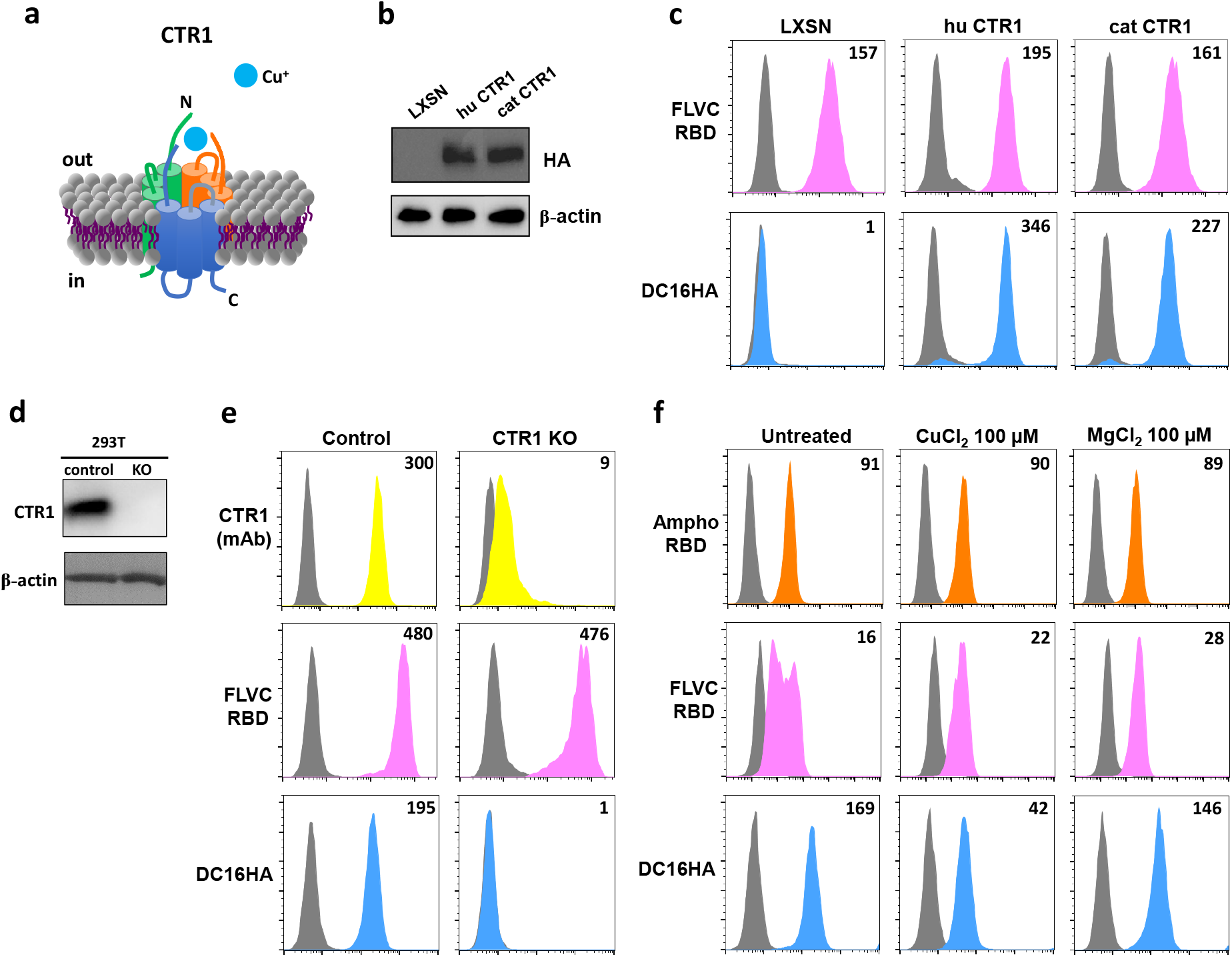
Refrex1 interacts with CTR1. **(a)** Graphical representation of the CTR1 copper transporter **(b)** Representative immunoblot of HA-tagged CTR1 in lysates from Chinese Hamster Ovary (CHO) cells stably transduced with MLV-based LXSN retroviral vector either empty or carrying the human (hu) or cat CTR1 cDNA. **(c)** CHO cells from (b) were evaluated for DC16HA and FLVC RBD binding by flow cytometry. Numbers indicate the specific change in mean fluorescence intensity of a representative experiment (n=3). **(d)** Detection of CTR1 by immunoblot in cell lysates of parental 293T or invalidated for CTR1 (CTR1 KO) generated by CRISPR/cas9 technology. **(e)** Cells from (d) were evaluated as in C using the indicated RBD ligands or an anti-CTR1 mAb. Representative experiment (n=3). **(f)** 293T cells grown in 100 µM CuCl_2_ or MgCl_2_ overnight were evaluated for CTR1 (DC16HA), PiT2 (Ampho RBD) and FLVCR cell surface expression by flow cytometry using indicated RBD ligands. Numbers indicate the specific change in mean fluorescence intensity of a representative experiment (n=3).

### Refrex1-CTR1 interaction regulates copper transport and homeostasis

Interaction between Retroviral Env and their cognate receptors often lead to decreased transporter activities ^23–26^. Since Refrex1 is derived from retroviral Env sequences, we tested whether its interaction with CTR1 would alter copper transport. We first expressed Refrex1 in 293T cells from ERV-DC7 and 16 *env* expression vectors either alone or in combination. Both vectors expressed ERV-DC7 and 16 *env* gene products with 3 copies of the HA tag at their C-terminus. We confirmed that both Refrex1 isoforms were properly expressed and secreted in culture medium of human 293T transfected cells (Fig. 4A) and that their presence alone or in combination conferred a strong resistance to infection by ERV-DC14 pseudotypes (Fig 4B), and abolished the binding of DC14RBD (Fig. 4C). We then measured the intracellular content of free copper in 293T cells by mass spectrometry. We found that the level of copper was reduced in the 3 conditions as compared to control, while levels of magnesium were unchanged (Fig 4D). We then performed the reverse experiment in feline cells, in which Refrex1 was silenced by transfection of siDC16. As expected, a specific increase in copper level was observed in the absence of Refrex1 in both FEA and CRFK cells (Fig 4E), suggesting that Refrex1 can control copper homeostasis by modulating copper transport through CTR1. We reasoned that changes in intracellular copper level would affect the expression of genes encoding actors of copper homeostasis like storage chaperones and transporters. Consistent with this hypothesis, we found an increase in *Atp7a* and to a lesser extent *Atp7b* gene expression both encoding copper exporters, as well as *Commd1* also assisting copper export (Fig 4F). Thus, Refrex1-depleted cells tend to reduce copper accumulation by expressing copper exporters and its partners. Surprisingly, *Ctr1* expression was not reduced upon Refrex1 depletion and copper accumulation, suggesting that feline cells were unable to regulate copper entry *via* the control of *Ctr1* transcription. We also found that Refrex1 depletion led to the up-regulation of *Sco1* and *Cox11* expression. Sco1 and Cox11 are two essential subunits of mitochondrial cytochrome c oxidase (COX-complex IV) that support the presence of copper ions for redox function. Their increased expression may be the sign of high electron transfer and oxidative phosphorylation (OXPHOS) activities. We found that absence of Refrex1 increased the level of reactive oxygen species (ROS) (Fig 4G) while ATP levels were poorly affected (Fig 4H). Surprisingly, we did not observe an increase neither in *Cox17* expression, a copper donor to Sco1 and Cox11 proteins, nor in *Sod1* expression encoding the copper/Zinc superoxide dismutase 1 that converts ROS into less toxic compounds. Overall, these results suggest that Refrex1 silencing leads to copper accumulation and subsequent reprogramming of gene expression involved in copper homeostasis and ROS production independent of ATP sysnthesis.

**Figure 4.**
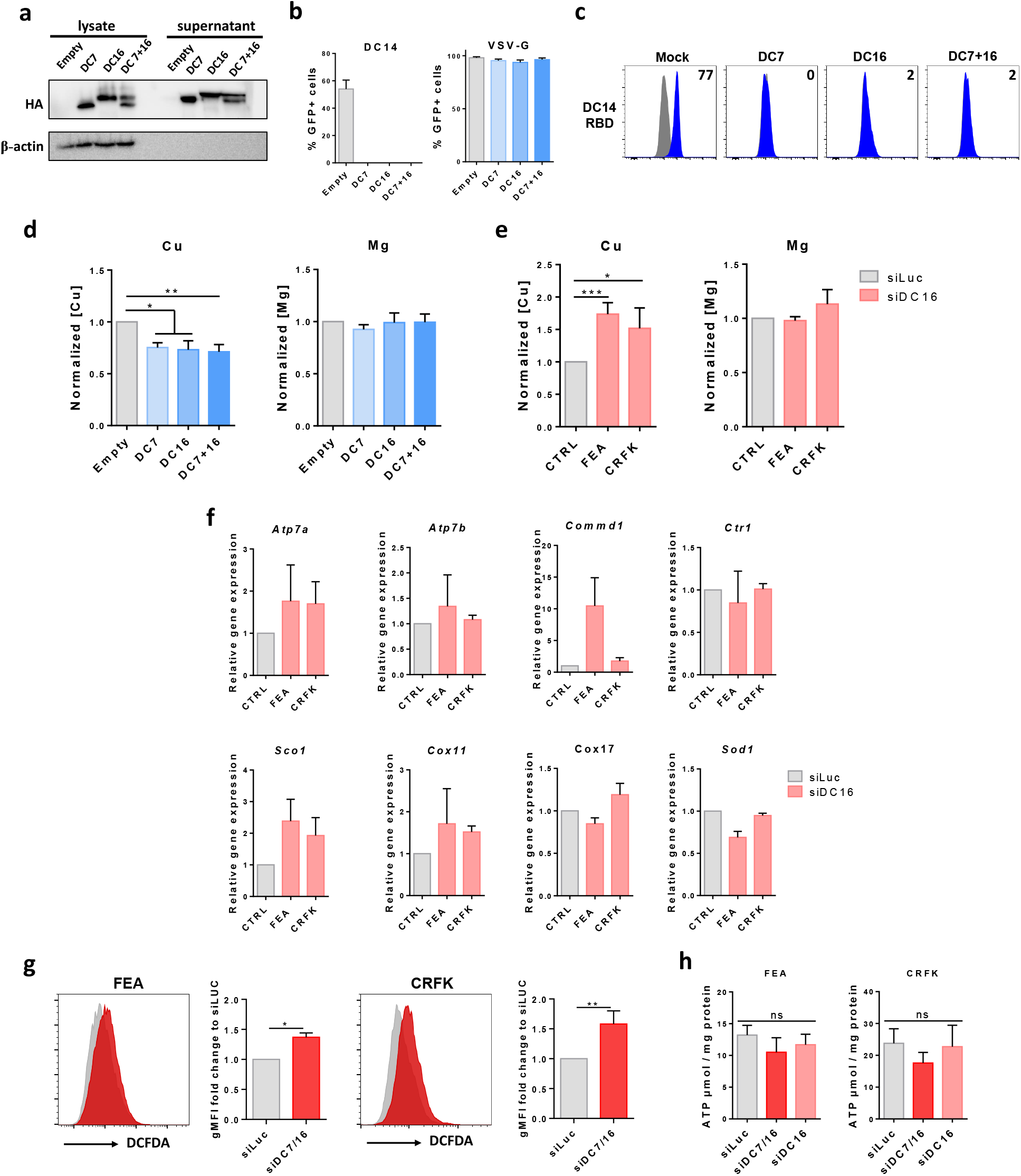
Altered copper homeostasis in the absence of Refrex1 in cat cells. **(a)** Detection of HA-tagged CTR1 by immunoblot in cell lysates and supernatants of 293T overexpressing empty vector pCHIX or DC7, DC16 alone or in combination (DC7 + DC16). **(b)** Sensitivity of cells from A to infection by EGFP lentiviral vectors pseudotyped with ERV-DC14 or VSV-G Env. Data are means ± SEM from n=3 experiments **(c)** Cells from (a) were evaluated for DC14RBD binding by flow cytometry. Numbers indicate the specific change in mean fluorescence intensity of a representative experiment (n=3). **(d)** Total amount of copper (Cu) or magnesium (Mg) were evaluated 48h post transfection by ICP-MS in 293T from A. One-way AVOVA with Dunnett’s multiple comparisons test * = p ≤ 0.05, ** = p ≤ 0.01. **(e)** Total amount of copper (Cu) or magnesium (Mg) were evaluated by ICP-MS 48h after transfecting feline FEA and CRFK with siLuc or siDC16. One-way AVOVA with Dunnett’s multiple comparisons test * = p ≤ 0.05, *** = p ≤ 0.001. **(f)** mRNA expression normalized to cat Gapdh of several genes involved in copper homeostasis (*Atp7a, Atp7b, Comm1, Ctr1, Sco1, Cox11, Cox17, and Sod1*) was evaluated by RT-qPCR in feline FEA and CRFK transfected with siDC16 or control siLuc. Values are expressed as the ratio of normalized expression to normalized expression in siLuc transfected cells. **(g)** Cellular reactive oxygen species (ROS) were detected using DCFDA staining in FEA and CRFK cells transfected with siDC7/16 or control siLuc. Data are mean ± SEM from at least n=4 experiments. Mann-Whitney test * = p ≤ 0.05, ** = p ≤ 0.01. **(h)** ATP quantification in FEA and CRFK transfected with either siDC16, siDC7/16 or a siLuc as a control. ATP assay was performed 24h post transfection. N=3 independent experiments. One-way AVOVA with Dunnett’s multiple comparisons test. ns = non-significant.

### Refrex1 expression is modulated by extracellular copper

Human cells can adapt to elevated copper concentration by reducing the level of CTR1 at the plasma membrane as seen in Fig 3F and as reported^3^. It is not known whether feline cells can sense and respond to extracellular copper in a similar manner. We attempted to detect CTR1 at the plasma membrane of feline cells, but neither our env ligand DC14RBD nor an anti-human CTR1 antibody were able to recognize the feline CTR1 by flow cytometry (Supplementary Fig 5), due to the capacity of Refrex1 to saturate CTR1. In contrast, CTR1 was weakly but reproducibly detected by flow cytometry using the DC16HA ligand (Fig 5A). We therefore evaluated CTR1 cell surface expression after exposure of FEA, CRFK and human cells to 100 µM of copper. As expected, we found that CTR1 was down-modulated in human 293T cells in the presence of elevated copper, but surprisingly enough, CTR1 expression on feline cells remained unchanged in the same conditions (Fig 5A). The same result was found at the transcriptional level (Fig 5B). Since feline cells have selected Refrex1 as a CTR1 cofactor and as a modulator of copper entry into cells, we next tested the possibility that feline cells would respond to an excess of extracellular copper by modulating Refrex1 expression. To test this hypothesis, FEA and CRFK cells were incubated with increasing amounts of copper and Refrex1 expression was assessed 24 hours later. We found that Refrex1 was up-regulated by elevated extracellular copper levels in a dose dependent manner (Fig 5C-D). Overall, these results suggest that adaptation of feline cells to copper excess does not involve CTR1 endocytosis as seen in human cells, but rather recruits Refrex1 and its CTR1 binding capacity to downmodulate copper acquisition.

**Figure 5.**
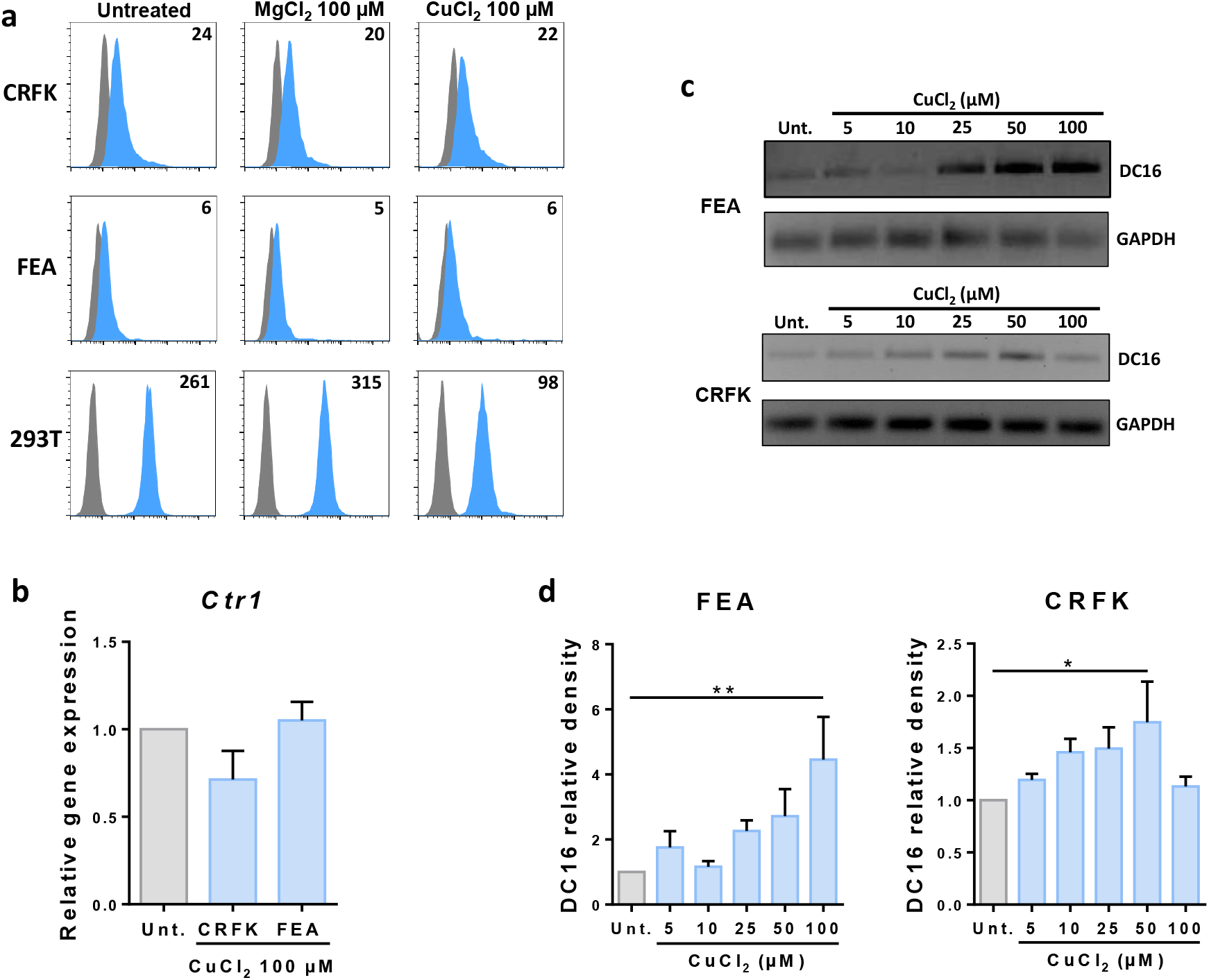
Refrex1 expression is modulated by extracellular copper level. **(a)** 293T and feline FEA and CRFK cells were incubated overnight with 100 µM CuCl_2_ or 100 µM MgCl_2_ and evaluated for CTR1 () cell surface expression by flow cytometry using DC16HA ligand. Numbers indicate the specific change in mean fluorescence intensity of a representative experiment (n=3). **(b)** Evaluation by RT-qPCR of *Ctr1* expression relative to Gapdh in feline FEA and CRFK cells incubated overnight with 100 µM copper. **(c)** Evaluation of DC16 and Gapdh expression by RT-PCR in feline FEA and CRFK incubated overnight with increasing concentrations of copper (5 to 100 µM). **(d)** Quantification of ERV-DC16 expression in cells from C expressed as the ratio to Gapdh expression. Data are means ± SEM from 3 experiments. One-way AVOVA with Dunnett’s multiple comparisons test p * = p ≤ 0.05 p ** = p ≤ 0.01

### Copper chelator BCS rescue Refrex1-silenced cells from death

The cell death observed in Refrex1-depleted feline cells correlates with the induction of ROS production and subsequent apoptosis, but the exact role of Refrex1 in this process is not known. Since Refrex1 can modulate copper entry *via* interaction with CTR1, we directly investigated the role of copper in cell death. FEA and CRFK cells transfected with siDC7/16 or control siLuc were grown for 72 hours in medium containing the copper chelator bathocuproinedisulfonic acid (BCS), and then cell survival was measured. We found that the induced cell death following Refrex1 silencing was reduced when extracellular copper was neutralized by BCS as observed by phase-contrast microscopy and crystal violet dosage for FEA cells and to a lesser extent for CRFK cells (Fig 6A and B). FEA cell rescue was dose-dependent since the percentage of cell survival gradually reached 100% of siLuc transfected cells with increasing amount of BCS. We next assessed the direct impact of copper chelation *via* BCS treatment on the induction of apoptosis following siDC7/16 transfection. At 24h, BCS decreased the proportion of annexin V-positive cells in FEA and, to a lesser extent, in CRFK cells (Fig 6C). Overall, these results suggest that, upon Refrex1 depletion by siRNA, an excess of copper could lead to an apoptotic cell death and that sensitivity to this copper-induced death is cell-type dependent.

**Figure 6.**
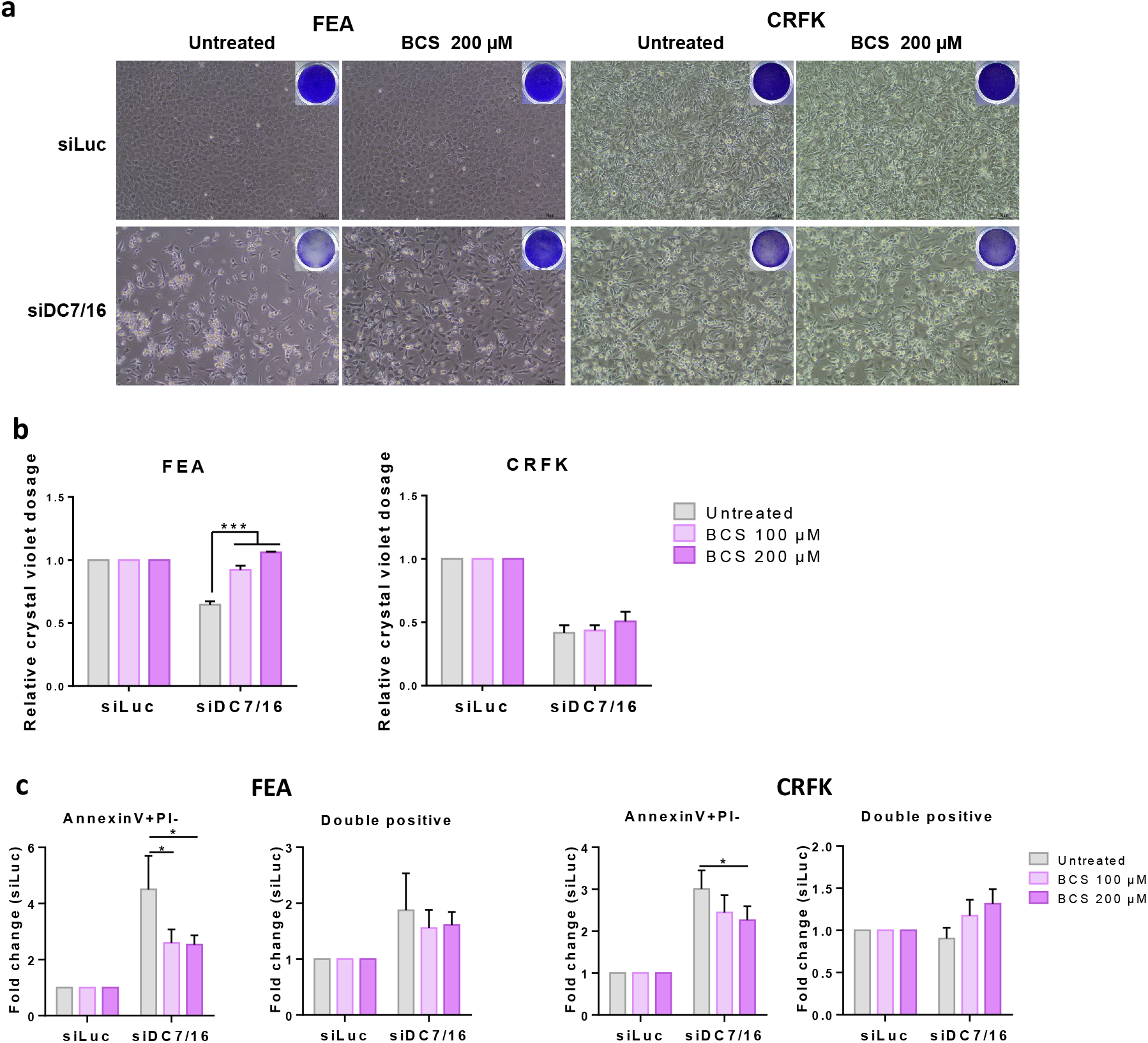
Copper chelator BCS protects Refrex1/DC16 -inactivated cells from death. FEA and CRFK cells were transfected with siDC7/16 or a siLuc control, and 5h post transfection medium was replaced with medium containing increasing doses of the copper chelator Bathocuproine disulphonate (BCS). **(a)** 3 days post-transfection bright light images were captured and corresponding wells were stained with crystal violet. Representative bright field images from the untreated and the 200μM BCS dose are shown and the insert shows the corresponding crystal violet image of the whole well. **(b)** Crystal violet of cells from (a) was solubilized and quantified. Data are mean ± SEM from n=3 experiments. One-way ANOVA with Dunnett’s multiple comparisons test *** = p ≤ 0.001. **(c)** At 24h post-transfection, cells were stained for Annexin V and Propidium Iodide (PI). The percentage of cells in the AnnexinV+/PI-as well as the double positive quadrants as defined in Fig 2 are shown. Data are mean ± SEM from n=4 or 5 experiments. Two-way ANOVA with Dunnett’s multiple comparisons test * = p ≤ 0.05.

### Presence of Refrex1 in cat serum

Presence of Refrex1 has been observed in the culture medium of all feline cell lines tested so far^16^, including in FEA and CRFK-CM (Fig 1D and Supplementary Fig 6). In contrast, Refrex1 is not expressed in all feline tissues, however expression of ERV-DC7 and 16 mRNA in peripheral blood mononuclear cells (PBMC) and other tissues may suggest that Refrex1 is present in cat serum^16^. To test this hypothesis, we measured the inhibitory activity of cat sera toward ERV-DC14 infection and DC14RBD binding. We found that cat serum strongly interfered with ERV-DC14 pseudotype infection and Env binding (Fig 7A and B) when 293T cells were preincubated with DMEM supplemented with cat serum. Serial dilutions of serum further revealed a 50% decrease in infection after a 500-fold dilution and a total loss of infection at a dilution threshold of 1000-fold, suggesting the presence of Refrex1 at high levels in this circulating body fluid, which may explain why cat red blood cells did not show any DC14RBD binding activity (Fig 7C). Finally, we tested whether the copper dependence of Refrex1 expression found in FEA and CRFK cells was also observed in cat PBMC. PBMC were isolated from blood of three domestic cat donors and incubated in the presence of copper for 24 hours. In the absence of copper, we found a high variability in Refrex1 expression from one donor to another and this expression was increased in presence of copper in PBMC of 2 cats out of 3 (Fig 7D). This suggest that copper sensing exists *in vivo* and that PBMC have the capacity to adapt their Refrex1 expression in response to copper variation in cat blood.

**Figure 7.**
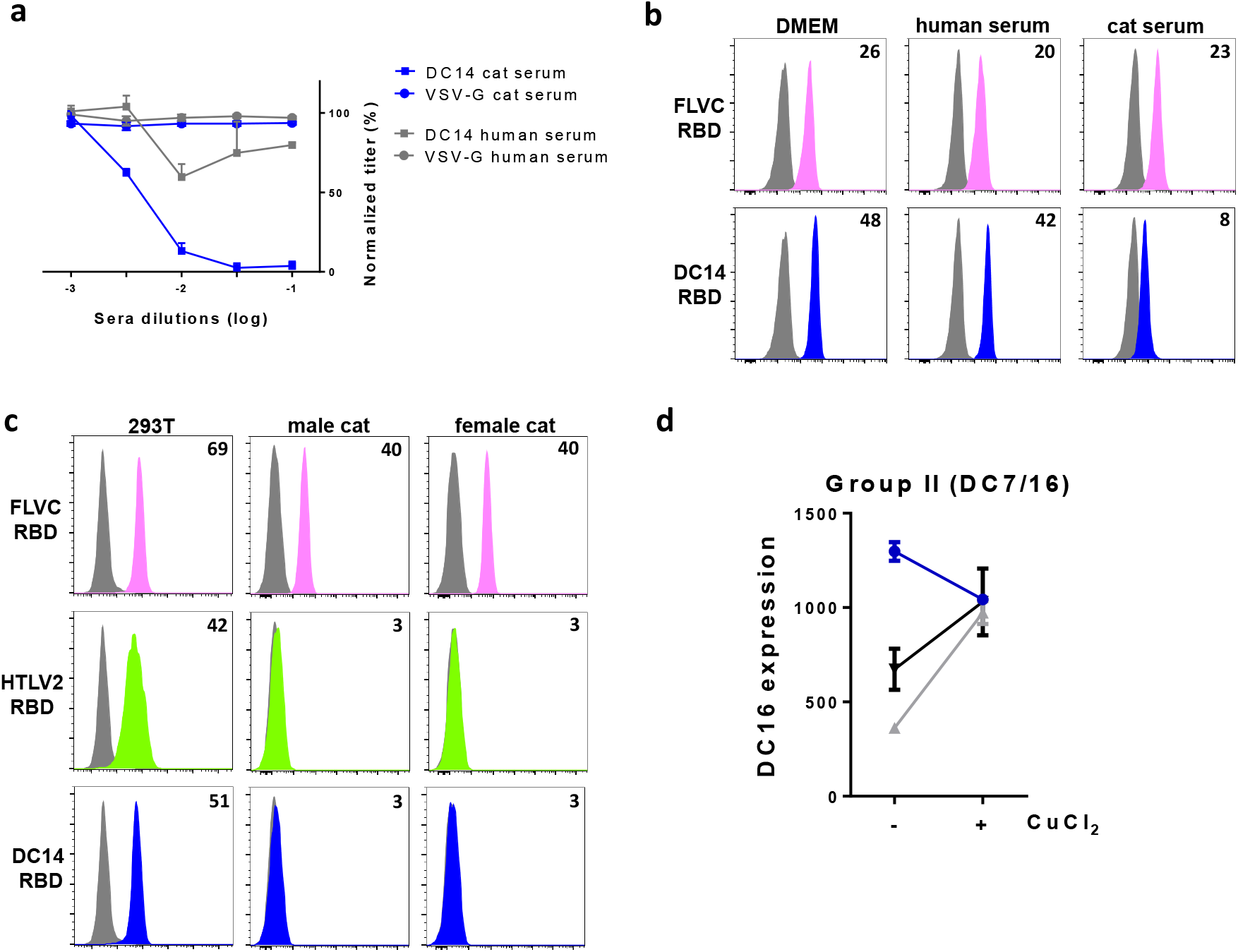
Refrex1 is present in cat serum. **(a)** 293T cells were incubated overnight with various dilutions of human and cat sera, and evaluated for their sensitivity to infection by EGFP lentiviral vector pseudotyped with ERV-DC14 or VSV-G Env. Data are means ± SEM from n=3 experiments. **(b)** Evaluation of cell surface expression of FLVCR and CTR1 on 293T cells incubated overnight with human and cat sera dilutions (1:50) by flow cytometry using the indicated RBD ligands. Numbers indicate the specific change in mean fluorescence intensity of a representative experiment (n=3). **(c)** Evaluation of cell surface expression of FLVCR, GLUT1 and CTR1 on human and cat red blood cells by flow cytometry using RBD ligands. Binding on 293T is shown as control. Numbers indicate the specific change in mean fluorescence intensity of a representative experiment (n=3). **(d)** Evaluation by RT-qPCR of ERV-DC16 *env* expression normalized with Gapdh in cat PBMCs isolated from whole blood of three independent cats and incubated overnight in 250 µM CuCl_2_.

## DISCUSSION

In this study, we describe a novel and unprecedented mechanism for copper homeostasis involving the regulation of CTR1-mediated copper acquisition by a co-opted retroviral Env, and we provide evidence that co-option of Refrex1, the Env product expressed from the ERV-DC7 and ERV-DC16 loci of modern cats, is crucial for cell survival. Refrex1 has maintained the capacity to interact with CTR1 during cat evolution and to block its retroviral receptor function from exogenous infection, explaining why Refrex1 was first identified as a restriction factor^16,32,37^. We further showed that interaction between Refrex1 and CTR1 impacted copper transport within cells and subsequent homeostasis as demonstrated by the lower level of copper in human cells overexpressing Refrex1. Conversely, downmodulation of Refrex1 expression in feline cells led to a significant increase in copper levels which in turn affected the expression of several genes involved in copper homeostasis. Surprisingly, Refrex1 behaves as an essential gene since we discovered that its downmodulation severely impaired cell survival, implicating that Refrex1 is a key regulator of both cellular copper level and toxicity. This copper-induced cell death was associated with significant increased ROS production and apoptosis levels, but further studies are required to uncover the underlying mechanisms. Thus, co-option of Refrex1 that occurred during cat evolution represents a novel layer of CTR1 regulation for the control of cellular copper homeostasis and redox fitness.

We observed a strong inhibitory activity against ERV-DC14 pseudotype infection in the serum of domestic cats, suggesting that plasma-resident Refrex1 can interact with cell surface CTR1 and this in nearly all cells of the cat organism. Red blood cells (RBC) are the major constituent of blood cells and they rely on a proper copper balance to control oxidative stress generated by high oxygen environment and by their heme iron-rich content. They are very sensitive to high copper levels, leading to ROS production, reduced antioxidant status and ultimately to hemolysis^38,39^. This suggests that bona-fide copper transport mechanisms exist on the surface of RBC, and consistent with this hypothesis, we detected the presence of CTR1 on human RBC using our DC14RBD ligand^35^. Surprisingly, CTR1 was not detected on feline RBC (Fig 7C), although feline CTR1 was clearly detected when overexpressed on CHO cells (Fig 3C). This may be the consequence of CTR1 saturation by Refrex1. It remains to be determined whether the interaction between CTR1 and soluble Refrex1 from blood can lead to copper transport modulation as reported with other soluble retroviral Env^25^.

In human cells, elevated copper levels trigger the endocytosis of CTR1 from the plasma membrane^3^ *via* the retromer-recycling function^40^, which prevent excessive copper uptake and toxic accumulation in cells. We found that this mechanistic adaptation does not occur in feline cells, but we discovered that CTR1 could be regulated in a different way with the help of Refrex1. In cells, Refrex1 is transported through the secretory pathway and is therefore not in contact with cytosolic copper, eliminating a role in sequestering or carrying copper just after entry *via* CTR1. On the contrary, Refrex1 is known to restrict virus entry through a mechanism known as interference to superinfection^16,41^, either by trapping CTR1 in the endoplasmic reticulum where they can interact or by saturating CTR1 at the plasma membrane with soluble Refrex1. Both possibilities lead to non-functional CTR1 but this does not explain how copper sensing may occur. Surprisingly, we found that Refrex1 expression was transcriptionally up-regulated in response to high levels of extracellular copper. At this point, we don’t know if this up-regulation resulted from the promoter activity of the ERV-DC7 or 16 long terminal repeat (LTR)^42^ or from an additional promoter of cellular origin. Interestingly, activation of HERV LTR by copper has already been reported, but the underlying mechanisms of copper action are not known^43^. Overall, our results suggest that cat cells can sense elevated copper levels and respond by stimulating Refrex1 expression and by downmodulating CTR1 transport activity through Refrex1-CTR1 interaction.

KO mouse models of CTR1 have shown an embryonic lethality suggesting that CTR1 defect is lethal in humans as well^44^. However, mutations in *CTR1* have not been associated with diseases. Nevertheless, copper-associated diseases have been reported, two of them involving copper pumps. Indeed, mutations in the *ATP7A* gene are associated with Menkes disease, a disorder of copper deficiency^45^, and mutations in the *ATP7B* gene cause Wilson disease, characterized by copper overload mainly in the liver, but also in other tissues such as brain^46^. In both cases, pathological mutations lead to copper unbalance with toxic consequences. We demonstrate here that Refrex1 is a key player of cellular copper homeostasis by regulating copper entry mediated by CTR1, and same as CTR1, ATP7A and ATP7B, Refrex1 integrity is crucial for cell physiology and survival. Copper disorders have been reported in several cats, some of which presented mutations in *Atp7b*^47–49^. Likewise, mutations in ERV-DC7 or ERV-DC16 affecting Refrex1 expression or interaction with Ctr1 may be associated with cat copper diseases, and as such *ERV-DC7/16* should be included in the list of candidate genes causing copper disorders.

As reported for Refrex1, the ability of ERV Env to be released in the extracellular environment and to modulate cell metabolism through receptor interaction could be widespread among vertebrates. Such a widespread co-option has been described for syncytins which play a crucial role in placentation in most mammals as well as marsupials^13^. In domesticated cats, FeLIX is another example of a soluble Env found in cat serum with receptor-binding capacity^50^. FeLIX has only been studied for its role as a facilitator of FeLV-T infection^27,51,52^, but its ability to interact with feline phosphate transporter Slc20a1 make it a potential modulator of phosphate influx and metabolism. Two other captured retroviral Env known to be secreted in the extracellular environment have been identified in humans. This includes the HERV-R Env expressed from the *erv3* locus that contains a stop codon before the transmembrane domain of the TM subunit^53^. This Env is thus non fusogenic but retains an intact SU with receptor-binding capacity, although 1% of Caucasian individuals have maintained a shorter Env looking like a RBD^54^. HEMO is another captured Env expressed from an ancestral *env* gene with an intact open reading frame^55^. Surprisingly, although HEMO is expressed as a SU-TM glycoprotein, it is shed in the extracellular environment and found at high level in the blood circulation of pregnant women^55^. Thus, the expression of soluble Env from co-opted ERV genes is not limited to Refrex1 and appears to exist in other organisms. Further studies are required to identify cellular partners of these soluble ERV factors in order to evaluate their role in physiological metabolic pathways and in metabolic disorders.

## MATERIALS AND METHODS

### Cell culture

HEK (Human Embryonic Kidney) 293T, hamster CHO (Chinese Hamster Ovary), and cat FEA (Feline embryonic fibroblast) and CRFK (Crandell feline kidney) cell lines were maintained in DMEM (Dulbecco’s Modified Eagle’s Medium, Gibco) supplemented with 10% fetal bovine serum (FBS, Sigma), 1% antibiotics (penicillin-streptomycin) and non-essential amino acids (Gibco). Cells were cultivated under humid atmosphere in a 5% CO_2_ incubator at 37°C. *CTR1* KO cells were generated by co-transfecting 293T cells with the pX458 vector carrying the Cas9 protease fused to GFP and a Sanger lentiviral CRISPR vector (Merck) carrying the BFP and a sgRNA against human CTR1 under the control of the U6 promoter (sgRNA: 5’-TACTAGCAATGTTCTATGAAGG-3’). GFP and BFP double positive cells were sorted 48 hour later using a BD FACSAria and cloned by limiting dilution in a 96-well plate.

### Plasmids, viral productions and infections

pLXSN vectors pseudotyped with the vesicular stomatitis virus G protein (VSV-G) were produced by cotransfecting 293T cells with the LXSN vectors carrying the human and feline CTR1 cDNAs^35^, the MLV Gag-Pol expression vector pC57GPBEB^56^ and the VSV-G Env expression vector pCSIG. Viral supernatants were harvested 48 hours after transfection, filtered through 0.45 µm pore-size filters and used to transduce CHO cell followed by G418 selection one day later. CSGW lentiviral vectors expressing the enhanced green fluorescent protein (EGFP) were produced by cotransfecting 293T cells with the pCSGW lentiviral vector, the HIV1 pSPAX2 *gag-pol* expression vector, and the Env expression vectors from VSV, ERV-DC14 or FeLV-C. They were used to infect for 48 hours 1×10^4^ cells/well seeded the day before in 96-well plate. Cells were then resuspended in 50 µl trypsin and 100 µl PBA (PBS with 2% FBS) and analyzed on a Novocyte flow cytometer (Acea, Biosciences, Inc). Data analyses measuring the percentage of EGFP-positive infected cells were performed using FlowJo software.

### Monitoring cell surface expression of retroviral receptors

Detection of CTR1, FLVCR, GLUT1 and PiT2 at the plasma membrane was performed using soluble receptor-binding domains (RBD) derived from ERV-DC14 (first 278 residues)^35^, FeLV-C (first 235 residues), HTLV2 and Amphotropic-MLV retrovirus fused to a mouse IgG1 Fc domain^24^. The Refrex1 ligand was generated by introducing 3 copies of the HA tag at the 3’ end of ERV-DC16 *env* in place of the stop codon. Human CTR1 was also detected using a mouse anti-CTR1 monoclonal antibody (Proteintech, 67221-1-Ig). RBD ligands were produced and used as previously described^57^. Briefly, 1×10^5^ cells were detached with trypsin 0.5%+1mM EDTA, resuspended in 100 μl PBA containing the different ligands and incubated at 37°C for 30 min. Cells were then washed twice with cold PBA and labeled for 20 min on ice with Alexa 488-conjugated anti-mouse IgG1 antibodies (1:400, ThermoFisher reference A21121). For the Refrex1/DC16HA ligand, cells were labeled beforehand for 30 minutes with an anti HA antibody (3F10, Roche 11867423001) diluted at 1:200 in PBA. For the mouse anti-CTR1 antibody, a 30 minute incubation was performed at a 1:200 dilution before the secondary antibody staining. Cells were then washed in PBA and analyzed on a Novocyte flow cytometer (Acea, Biosciences, Inc). Data analysis was performed using FlowJo software. When indicated, 293T were incubated in FEA or CRFK cells conditioned medium, or incubated overnight with 100 µM CuCl_2_ (307483, Sigma) or MgCl_2_ (208367, Sigma).

### RNA isolation, synthesis of cDNA, RT-PCR and RT-qPCR

Total RNA (5×10^6^ cells) was extracted from FEA or CRFK cell lines by using the RNeasy Plus Mini Kit (Qiagen). RNA from FEA, CRFK cells or cat PBMCs were primed with Oligo(dT)18 and cDNA were synthetized using the Superscript II First Strand Synthesis System (Invitrogen). A 523bp fragment of ERV-DC7 or ERV-DC16 env genes was then amplified by PCR using One Taq DNA Polymerase (New England Biolabs) with the following primers: DC7/16-Fwd (5’-AATTGGAGCTAAAAATGTCGGG-3’) and DC7/16-Rev (5’-TTCCCAATAATGATACCCTTC-3’). The PCR products were agarose gel purified using the PCR clean-up Gel Extraction kit (Macherey Nagel) according to manufacturer’s protocol and used for direct Sanger sequencing. For RT-qPCR, cells were lysed and RNA extracted using the Luna Cell Ready Lysis Module (E3032S, New England Biolabs) according to the manufacturer’s protocol. RT-qPCR on extracted RNA was performed using the Luna One-Step RT-qPCR Kit (M300, New England Biolabs). The samples were analyzed using a LightCycler 480 instrument (Roche). qPCR primers for ERV-DC *env* genotypes I, II, III have been previously described as follow:

Genotype I, Fwd 5’-GCTTGCACTTCCACCAGTTG-3’ and rev 5’-ACCTGTTCCTGTCTTGCGTAG-3’;

genotype II, Fwd 5’-ACCTGTTCCTGTCTTGCGTAG-3’ and rev 5’-TGCCAACTGGTTTTGTTACTTATG-3’;

Genotype III, Fwd 5’-GCCTCCCTACCCGACTTCC-3’ and rev 5’-AGGGGGTTTAGCCGTTAGG-3’.

qPCR primers for feline genes are as follow:

Atp7a, Fwd 5’-GCTGTGTGGCTTGATTGCTA-3’ and rev 5’-TTTCCCCTCCTCCTGAAGTT-3’;

Atp7b, Fwd 5’-CTGTAAAGAGGAGCTGGGAAC-3’ and rev 5’-CCTCCACGTTGCTGACTTTA-3’;

Commd1, Fwd 5’-CATGTGACCAAGCTGCTGTT-3’ and rev 5’-CAGGCAACCCTGACTTGTTT-3’ ;

Sco1, Fwd 5’-GGAATTCCAACTCTGCCAAA-3’ and rev 5’-CTGGGGCCAGGACTGTAATA-3’ ;

Cox11, Fwd 5’-ATCGGCTGTATTGCCAGACT-3’ and rev 5’-AAAACGCCAGTGCAGTCTCT-3’ ;

Cox17, Fwd 5’-GGAGAAGAAGCCTCTGAAGCC-3’ and rev 5’-TGTGGGCCTCAATTAGGTGTC-3’;

Ctr1, Fwd 5’-GGAACCATCCTTATGGAGACAC-3’ rev 5’-GCTGATGACCACTTGGATGATA -3’;

Sod1, Fwd 5’-TGCACTCTCAGGAGACCATTCC-3’ and rev 5’-CCACAAGCCAAACGACTCCCAG-3’;

Gapdh, Fwd 5’-GTCTTCCTGCGACTTTAACAGTG-3’ and rev 5’-ACCACCCGGTTGCTGTAGCCAA-3’.

### Immunoblotting

Cells were lysed in 50 mM Tris-HCl (pH 8.0), 100 mM NaCl, 1 mM MgCl_2_, 1% Triton X-100, and protease inhibitors (Complete, Roche). Cell extracts were separated on a 12.5 % SDS acrylamide gel, transferred to PVDF membranes and protein expression was detected using anti HA (3F10, Roche 11867423001, 1:5000) followed by horseradish peroxidase (HRP)-conjugated anti-rat antibodies (Sigma, A9037, 1:10000) or CTR1 monoclonal antibody (Proteintech, 67221-1, 1:1000) followed by horseradish peroxidase (HRP)-conjugated anti-mouse antibodies (Jackson, 715-035-150, 1:10000) or anti ß-actin-HRP (A3854, Sigma, 1:100000) antibodies. Signals were visualized using the Luminata Forte detection reagent (Merck Millipore) and recorded using the Chemidoc gel imaging system.

### Crystal violet viability staining

FEA and CRFK cells were seeded in 24-well plates at a density of 1×10^5^ and 8×10^4^ cells per well respectively. The next day, they were transfected with siDC7/16 or siLuc as a control using JetPrime according to the manufacturer’s protocol. 6h later, the medium was changed to medium without BCS or containing increasing concentrations of BCS (100 µM, 200 µM). 3 days post-siRNA transfection, pictures of each well were taken using a Nikon Eclipse Ts2 microscope. Cells were then washed with PBS and fixed with formalin (Sigma HT5011) for 20min at room temperature. Formalin was removed before adding Crystal violet 0.5% in 20% methanol and incubating a minimum of 1h at room temperature. Crystal violet was then washed 3 to 4 times in water before being left to dry. Once dry, the plate was scanned to get an image of the overall density of cells in all wells. Crystal violet was then solubilized in 1 mL 80% methanol and 100 µL of solution were distributed in 3 replicate wells of a 96-well plate. Absorbance at 570 nm was read using a Infinite F200PRO microplate reader (Tecan).

### Copper dosage

FEA, CRFK and 293T cells were seeded in 6-well plates at a density of 5×10^5^ per well. The next day, they were transfected using JetPrime according to the manufacturer’s protocol with siDC16 or siLuc as a control for FEA and CRFK or with DC7, DC16, DC7+16 or pCHIX as a control for 293T cells. The medium was changed 4h later and replace by a medium with 100 µM BCS. The next day, the medium was removed and cells were incubated 1h with 25 µM CuCl_2_. Then they were washed twice with cold PBS and 200 µl lysis buffer (Thermofisher, 87788) supplemented with protease inhibitors (complete, Roche) was added in each well. A portion (5 µl) was kept to quantify the total amount of proteins using Pierce™ BCA protein assay kit (23225, ThermoFisher). 5 ml of HNO3 5% (Sigma, 84378) were added to each sample and they were placed 1h at 95°C. Copper and magnesium dosage were then quantified using the inductively coupled plasma mass spectrometry (ICP-MS) method.

### Viability assay

FEA and CRFK cells were seeded in 6-well plates at a density of 3×10^5^ cells per well. The next day, they were transfected with siDC7/16, DC16 or siLuc or siGPX4 as controls using JetPrime according to the manufacturer’s protocol. 6h later, cells were harvested using trypsin 0.5% + 1 mM EDTA and then distributed into 6 technical replicates in a 96-well plate. Cell viability was measured 48h later (or otherwise indicated) using CellTiter 96® AQueous One Solution Cell Proliferation Assay (G3582, Promega) according to the manufacturer’s instructions. Absorbance was measured at 490 nm using an Infinite F200PRO microplate reader (Tecan).

### ATP measurement

FEA and CRFK cells were seeded in 96-well plates at a density of 3×10^4^ cells per well. The next day, they were transfected with siDC7/16, siDC16 or siLuc or siGPX4 as controls using JetPrime according to the manufacturer’s protocol. ATP measurement was performed using the ATPlite Luminescence Assay System (6016941, PerkinElmer) according to the manufacturer’s protocol.

### Annexin V/Propidium Iodide staining

FEA and CRFK cells were seeded in 6-well plates at a density of 3×10^5^ and 2.5×10^5^ cells per well respectively. The next day, they were transfected with siDC7/16, siDC16 or siLuc as a control using JetPrime according to the manufacturer’s protocol. The medium was changed 6h later. 24h post-siRNA transfection, cells were washed with PBS and detached using Trypsin 0.5%+1mM EDTA for 5min at 37°C. Cells were collected using complete medium and distributed into 3 technical replicates in a 96-well plate. Cells were then washed with PBS and stained for Annexin V and propidium iodide (PI) using the eBioscience Annexin V-FITC Apoptosis detection kit (cat# BMS500FI) according to manufacturer’s instructions with slight modifications. Briefly, cells were washed once in 1X binding buffer then Annexin V was diluted 1:40 in binding buffer and 40 µL was added per well. Cells were resuspended and incubated for 15 min at room temperature in the dark. After one wash in binding buffer, cells were resuspended in 150 µL of binding buffer containing PI diluted 1:20 and directly analyzed using a Novocyte flow cytometer (Acea, Biosciences, Inc). Data analysis was performed using FlowJo software. One well was not stained with Annexin V and stained with PI only to be used as a gating control.

### ROS staining

Cells were transfected with siRNA, incubated and collected as described above. Cells were then distributed into 3 or 4 technical replicates in a round-bottom 96-well plate and washed twice with PBS. Cells were resuspended in 50 µL of PBS containing 2 µM of DCFDA and incubated at 37°C in the incubator for 30 min to allow for dye loading. After 1 wash in PBS, cells were resuspended in 50 µL PBS+SYTOX Red diluted 1:1000 and incubated for 15 min at room temperature in the dark. 100 µL of PBS was then added per well and cells resuspended before analysis using a Novocyte flow cytometer (Acea, Biosciences, Inc). Data analysis was performed using FlowJo software. One well was not stained with DCFDA to be used as a control of background cell fluorescence.

### PBMCs purification

The isolation of peripheral blood mononuclear cells (PBMCs) from whole blood of cats (1 mL taken in EDTA tubes) given by veterinary clinics with the consent of cat owners was performed using Ficoll (Histopaque, 11191, Sigma). Briefly, total sample of whole blood of each cat was deposited on 2mL of the density gradient and centrifuge at 600g during 30min at room temperature without brake. The ring of PBMC was carefully recovered and wash twice in PBS. The cells were then resuspended in RPMI (Roswell Park Memorial Institute)-1640 medium supplemented with 10% fetal bovine serum (FBS, Sigma), 1% antibiotics (penicillin-streptomycin) and when indicated, were incubated overnight with 250 µM CuCl_2_.

### Statistical analysis

Statistical analyses were performed using GraphPad Prism 6 software. The data were expressed as the mean ± SEM. The results were considered statistically significant at a p-value <0.05 (*), <0.01 (**), <0.001 (***) or <0.0001 (****). Statistical tests and number of independent experiments are indicated in the figure legends.

## ACKNOWLEDGEMENTS

The authors are grateful to the Centre Hospitalier Vétérinaire Languedocia, the clinique vétérinaire de l’Aiguelongue, the clinique vétérinaire du Corum, and the clinique vétérinaire d’Alco in Montpellier France for providing blood samples, to the OSU OREME platform of Montpellier University for metal dosage, and to Andrea Cimarelli (CIRI, Lyon, France) for CRFK cells. We also thank all the members of our team for technical help and advice. This work was supported by the French National Research agency (ANR CALCIPHOS, ANR-17-CE14-0008-01 to JLB) and by the French SIDACTION (19-2-AEQ-12560 to JLB). ST was supported by the ANR CALCIPHOS, LC by the French AIDS National Research ANRS, VC by CNRS and JLB by INSERM.

## AUTHOR CONTRIBUTIONS

ST, LC, VC and JLB conceived, designed and performed experiments, and analyzed the data. JLB wrote the original draft. ST, LC, VC and JLB edited the manuscript.

