## Supplemental for "A co-opted endogenous retroviral envelope promotes cell survival by controlling SLC31A1/CTR1-mediated copper transport and homeostasis"

**SUPPLEMENTARY FIGURES AND LEGENDS**

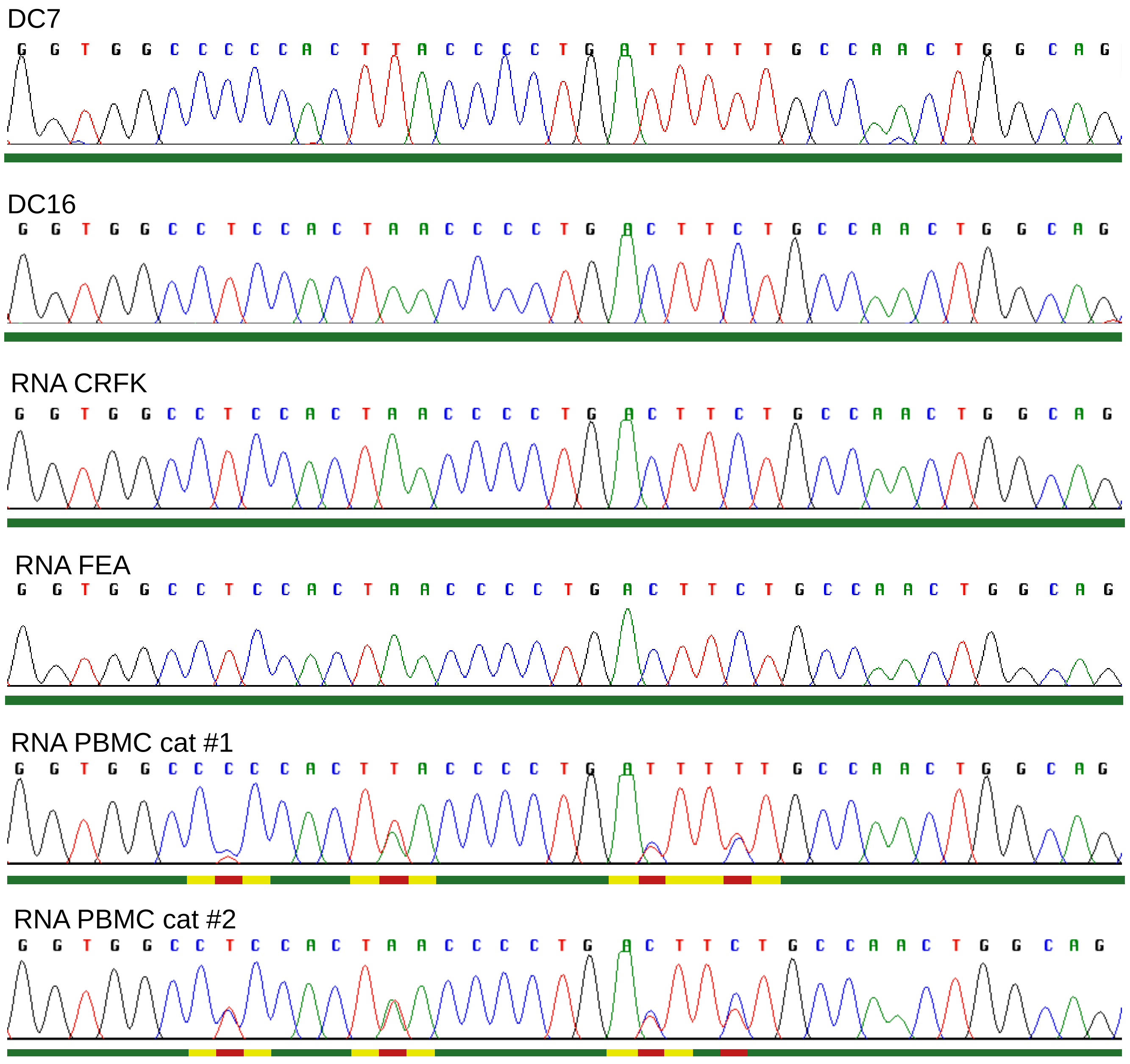


**Supplementary figure 1: Direct DNA Sanger sequencing chromatograms showing sequence polymorphisms of Refrex1 (DC7 or DC16) in cDNA from FEA, CRFK cells or cat PBMCs**

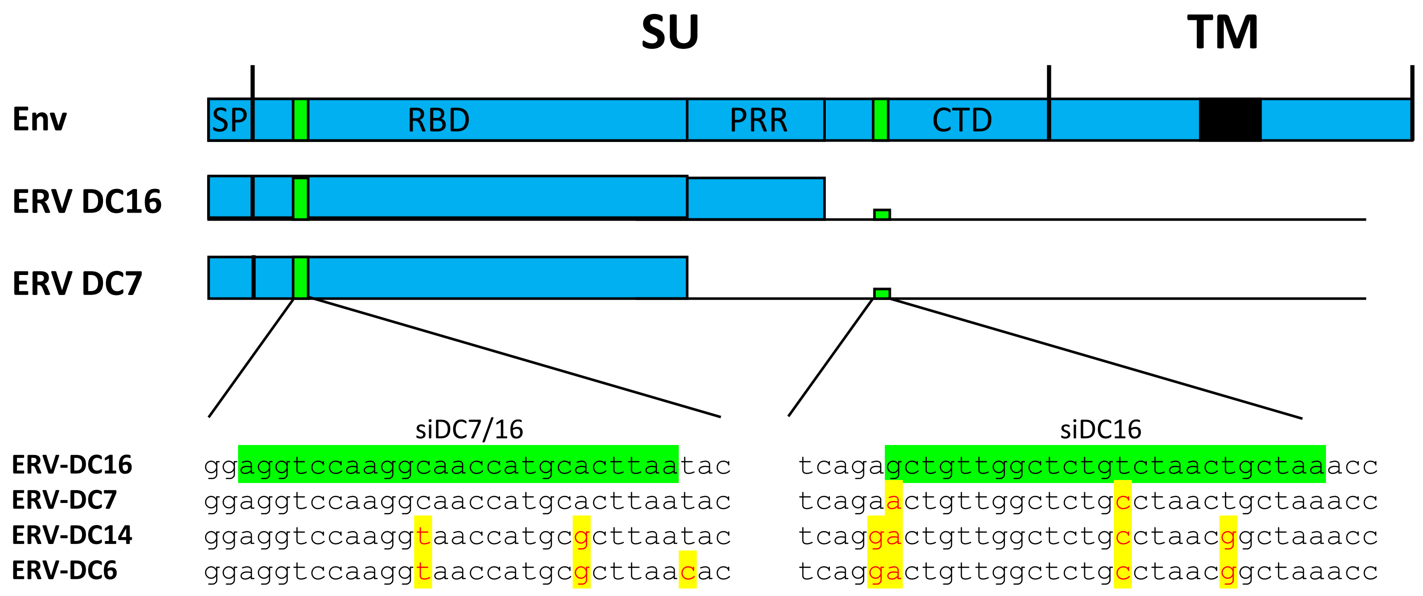


**Supplementary figure 2: Sequences of siDC7/16 and siDC16**

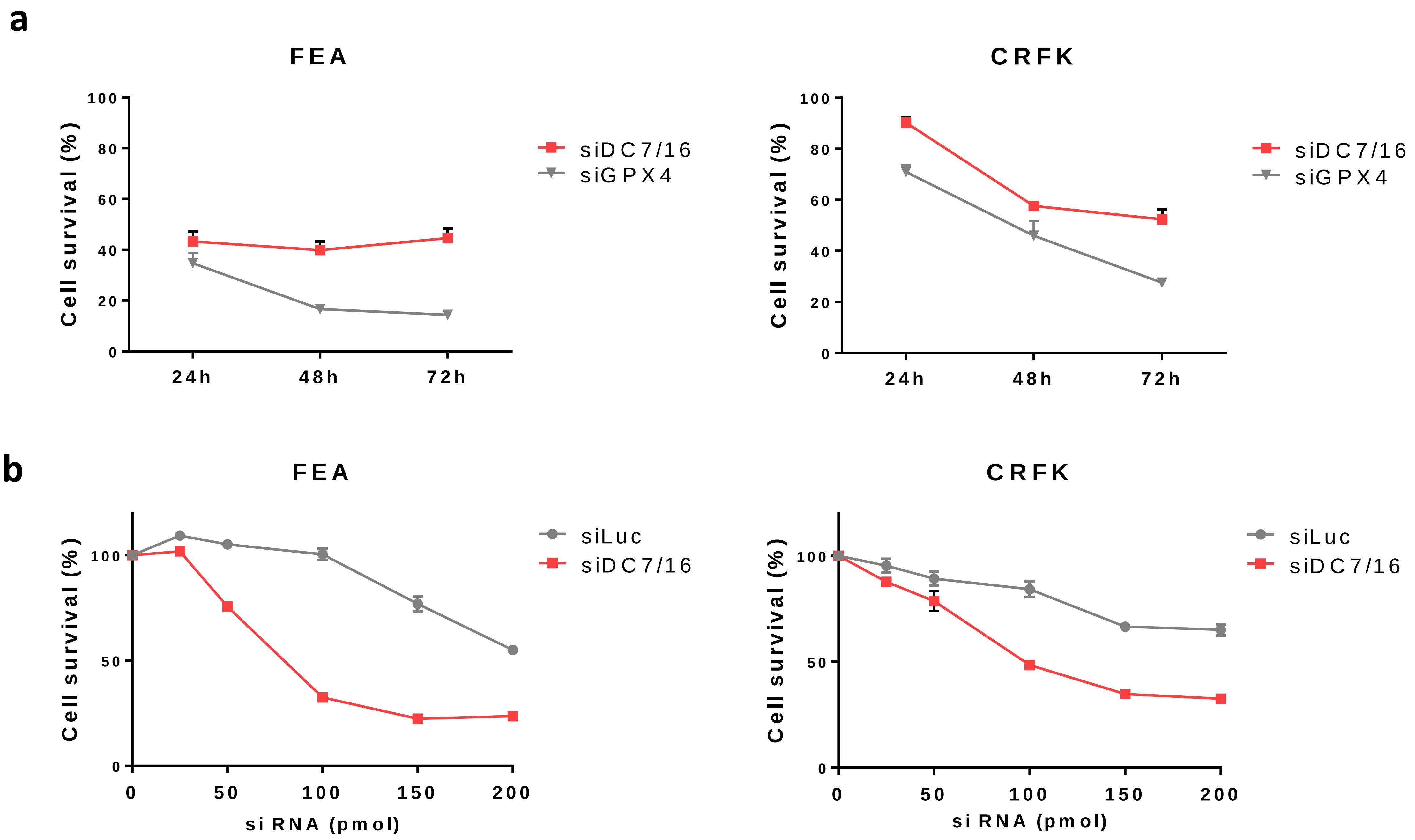


**Supplementary figure 3: FEA and CRFK cell toxicity following siDC7/16 transfection is time- and dose- dependent**

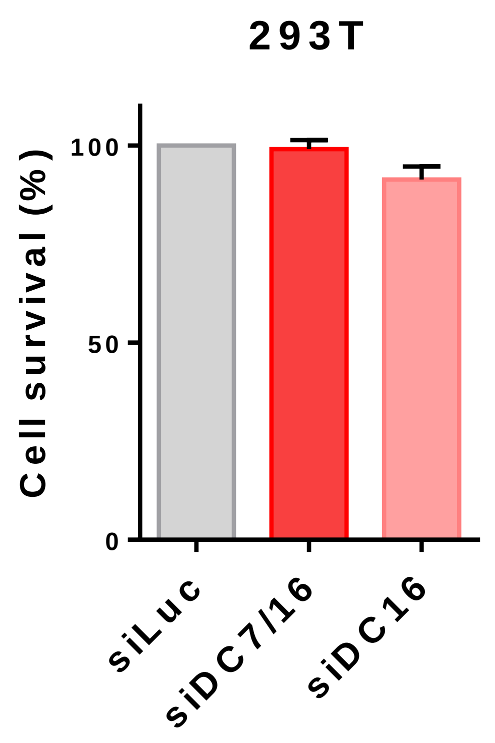


**Supplementary figure 4: Human 293T cells are not sensitive to siDC7/16 and siDC16 transfection**

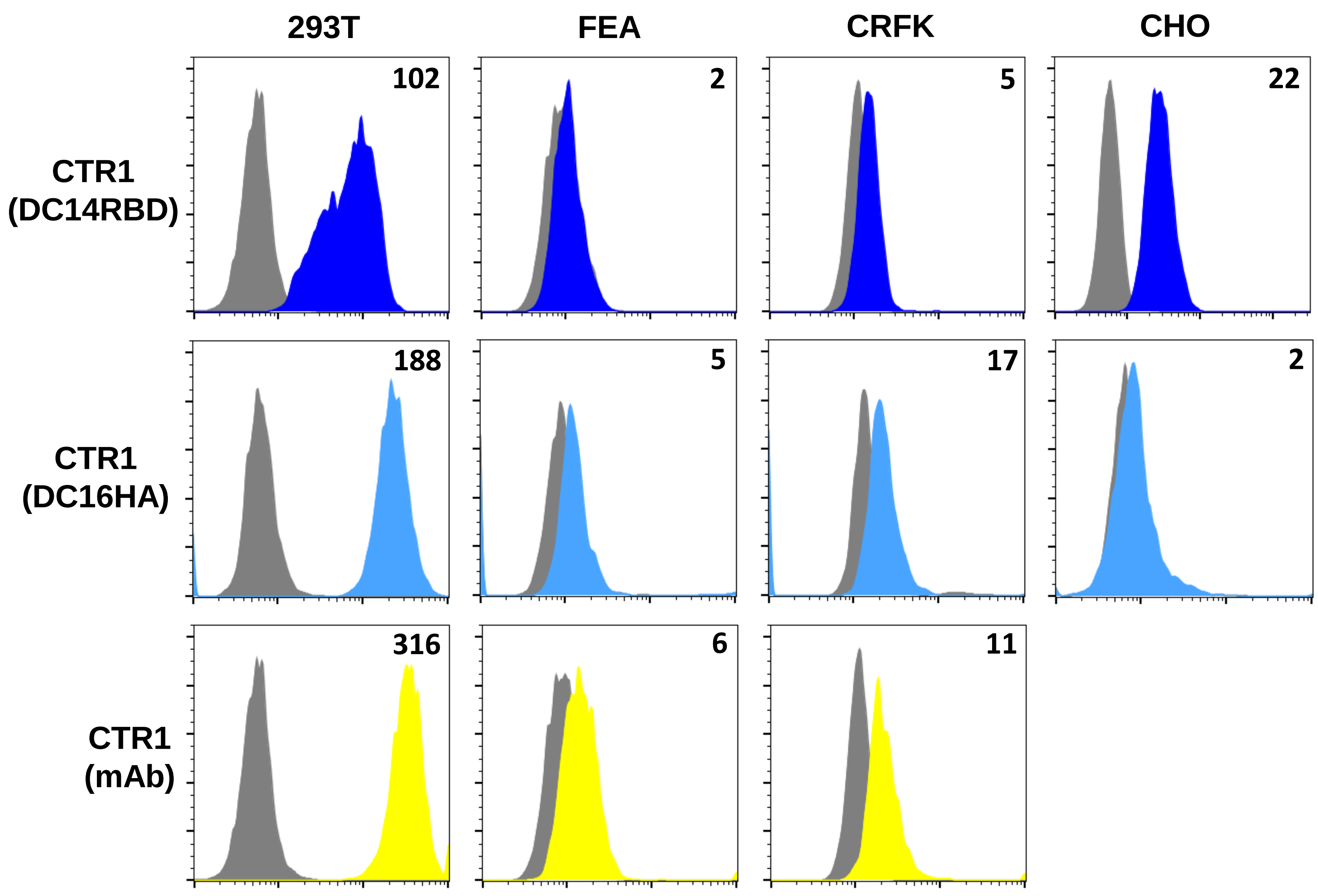


**Supplementary figure 5: Rabbit anti–CTR1 antibody and DC14RBD are unable to detect CTR1 at the surface of feline cells**

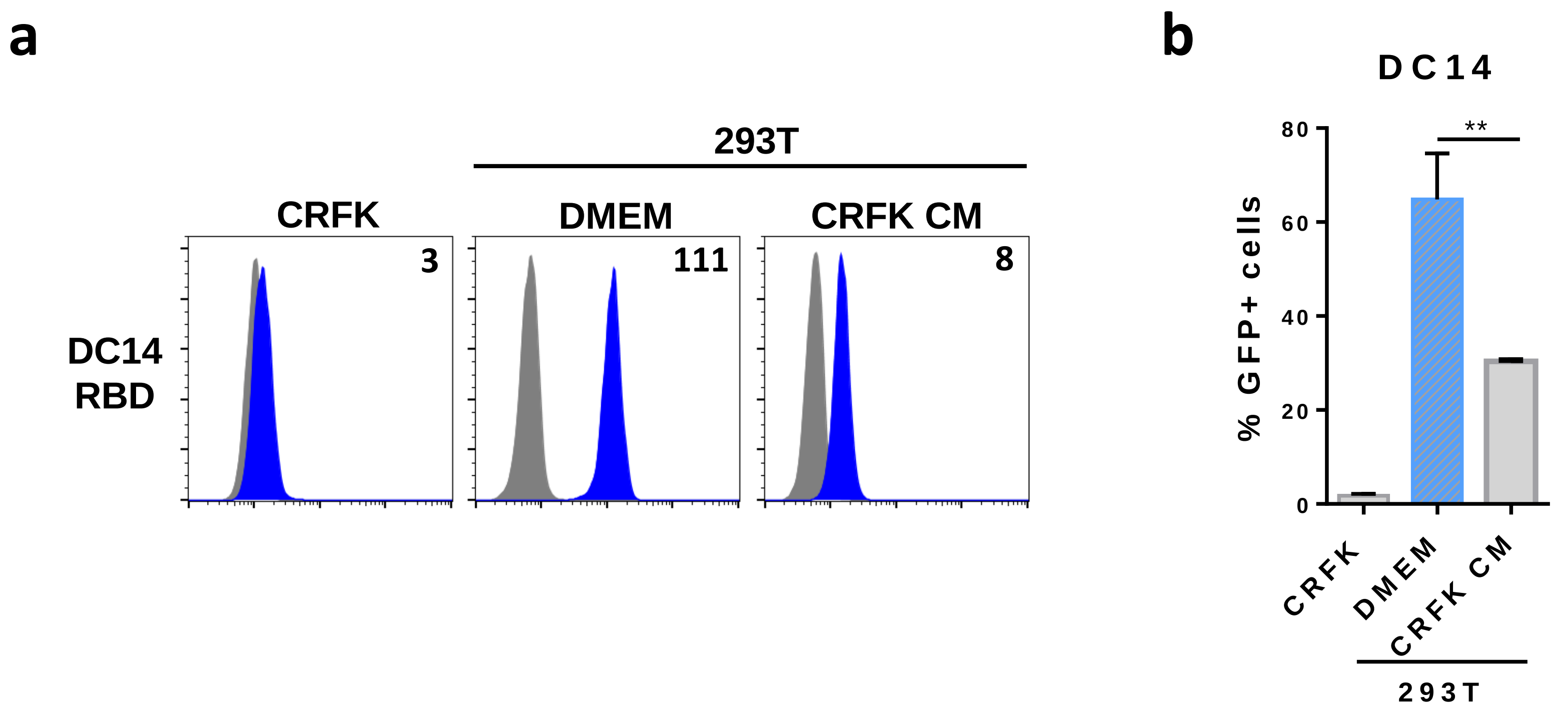


**Supplementary figure 6 : Refrex1 is present in CRFK CM**

***Supplementary figure 1: Direct DNA Sanger sequencing chromatograms showing sequence polymorphisms of a fragment of Refrex1 (DC7 or DC16) in cDNA from FEA, CRFK cells or cat PBMCs.***

Total RNA were extracted from the indicated cells and sanger sequencing was performed on RT-PCR amplified cDNA. A 39-bp DNA fragment starting at position 178 from start codon and containing 4 differences between ERV-DC7 and ERV-DC16 env is shown.

***Supplementary figure 2: Sequences of siDC7/16 andsiDC16.***

Schematic representation of ERV-DC7 and ERV-DC-16 Env organization. Both Env contain a signal peptide (SP) and a receptor-binding domain (RBD). Only DC16 contains an extra proline-rich region (PRR). SU: surface unit; TM: transmembrane unit; CTD: C-terminal domain. Partial sequence alignments comprising the siDC7/16 and siDC16 sequences are shown. ERV-DC7 and 16 are prototypes of genotype group II, ERV-DC14 of genotype group I and ERV-DC6 of genotype group III.

***Supplementary figure 3: FEA and CRFK cell toxicity following siDC7/16 transfection is time- and dose-dependent***

**(a)** Feline FEA and CRFK transfected with a siDC7/16 or siLuc control were evaluated for cellular viability 24h, 48h or 72h post transfection using CellTiter 96® AQueous One Solution Cell Proliferation Assay. **(b)** Feline FEA and CRFK were transfected with increasing quantities (25-200 pmol) of siDC7/16 or siLuc and evaluated for cellular viability 48h post transfection using CellTiter 96® AQueous One Solution Cell Proliferation Assay.

***Supplementary figure 4: Human 293T cells are not sensitive to siDC7/16 and siDC16 transfection***

Human 293T transfected either with siLuc, siDC7/16 or siDC16 were evaluated for cellular viability 48h post transfection using CellTiter 96® AQueous One Solution Cell Proliferation Assay.

***Supplementary figure 5: Mouse anti–CTR1 monoclonal antibody and DC14RBD are unable to detect CTR1 at the surface of feline cells***

293T and feline FEA and CRFK cells were evaluated for CTR1 cell surface expression by flow cytometry using the DC14RBD and DC16HA ligands or an anti-CTR1 antibody (mouse monoclonal 1:1000). Numbers indicate the specific change in mean fluorescence intensity of a representative experiment (n=3).

***Supplementary figure 6: Refrex1/DC16 is present in CRFK CM***

**(a)** Detection of DC14RBD binding was performed on CRFK cells as well as on 293T cells pre-incubated for 5 hours with either DMEM or CM from CRFK cells by flow cytometry. Numbers indicate the specific change in mean fluorescence intensity of a representative experiment (n=3). **(b)** Cells treated as in A were evaluated for their sensitivity to infection by EGFP lentiviral vector pseudotyped with ERV-DC14. Data are means ± SEM from n=3 experiments. Student unpaired t-test, **=p≤0.01.
